# Flagellin from *Pseudomonas aeruginosa* modulates SARS-CoV-2 infectivity in CF airway epithelial cells by increasing TMPRSS2 expression

**DOI:** 10.1101/2020.08.24.264564

**Authors:** Manon Ruffin, Jeanne Bigot, Claire Calmel, Julia Mercier, Andrés Pizzorno, Manuel Rosa-Calatrava, Harriet Corvol, Viviane Balloy, Olivier Terrier, Loïc Guillot

## Abstract

The major challenge of the COVID-19 health crisis is to identify the factors of susceptibility to SARS-Cov2 in order to adapt the recommendations to the populations and to reduce the risk of getting COVID-19 to the most vulnerable people especially those having chronic respiratory diseases including cystic fibrosis (CF). Airway epithelial cells (AEC) are playing a critical role in the immune response and in COVID-19 severity. SARS-CoV-2 infects the airways through ACE2 receptor and the host protease TMPRSS2 was shown to play a major role in SARS-CoV-2 infectivity. Here, we show that the main component of *P. aeruginosa* flagella, ie. flagellin is able to increase TMPRSS2 expression in AEC, and even more in those deficient for CFTR. Importantly, this increased TMPRSS2 expression is associated with an increase in the level of SARS-CoV-2 infection. Considering the urgency of the health situation, this result is of major significance for patients with CF which are frequently infected and colonized by *P. aeruginosa* during the course of the disease.

---

As of November 3, 2020, coronavirus disease 2019 (COVID-19) pandemic caused by the Severe Acute Respiratory Syndrome (SARS)-Coronavirus (CoV)-2 has infected globally near 47 million people with more than 1 million deaths (https://covid19.who.int). One of the major challenges of this health crisis is to identify factors of susceptibility to this viral infection in order to adapt public health recommendations and to reduce the risk of getting COVID-19, notably for the most vulnerable people. This is the case of patients having common chronic respiratory diseases (CRD) such as asthma and chronic obstructive pulmonary disease (COPD), but also for patients with less common or rare CRD, such as cystic fibrosis (CF) and interstitial lung diseases. Even if it seems reasonable to think that, as they have a lung impairment, people with CRD would be at higher risk to develop a severe COVID-19, the magnitude of this risk is still uncertain (1). Along with the clinical follow-up of these patients to have a better estimation of their risk, basic research regarding the pathophysiology of SARS-CoV-2 infection will provide important insight.

This is particularly true for patients with CF. This disease is caused by variants in the *CFTR* gene, the most frequent being F508del, which result in an abnormal function of the airway epithelial cells (AEC). During the course of CF, lungs of the patients are inflamed and chronically infected by various pathogens including *Pseudomonas aeruginosa,* which is the most prevalent (80% in CF adults) (2). Up to now, a multinational report identified 40 cases of CF patients infected by SARS-CoV-2 with no death recorded (3). Of these cases, 50% had chronic *P. aeruginosa* airways infection, 78% had symptoms at COVID-19 onset such as fever (77% of the symptomatic patients), altered cough (48%) and/or dyspnea (32%). AEC are known to play a critical role in both the immune response and COVID-19 severity (4). SARS-CoV-2 infects the airways mainly through the ACE2 receptor, and two specific host proteases, TMPRSS2 and FURIN, were shown to play a major role in SARS-CoV-2 infectivity (5–9).

In this study, we show that the main component of *P. aeruginosa* flagella, ie. flagellin (*PA*-F), is able to increase TMPRSS2 expression in airway epithelial cells, and even more in those deficient for *CFTR*. Importantly, this increased TMPRSS2 expression is associated with an increase in the level of SARS-CoV-2 infection.

We first examined *ACE2*, *FURIN* and *TMPRSS2* expression from a previous transcriptomic study performed with primary human airway epithelial cells (hAEC) isolated from control donors (non-CF) and CF patients homozygous for the *CFTR* F508del variant, infected by *P. aeruginosa* (10). At baseline (time 0 h), similar *ACE2* and *FURIN* mRNA expression levels are observed between non-CF and CF primary hAEC (**Fig. 1A, left**), whereas *TMPRSS2* expression is significantly increased in CF primary hAEC (**Fig. 1A, right**). Importantly, *P. aeruginosa* increases *TMPRSS2* mRNA expression over time (**Fig. 1A**) in CF primary hAEC only, while *ACE2* and *FURIN* expressions are not affected (**Fig. 1A, left**).

**Figure. 1.**
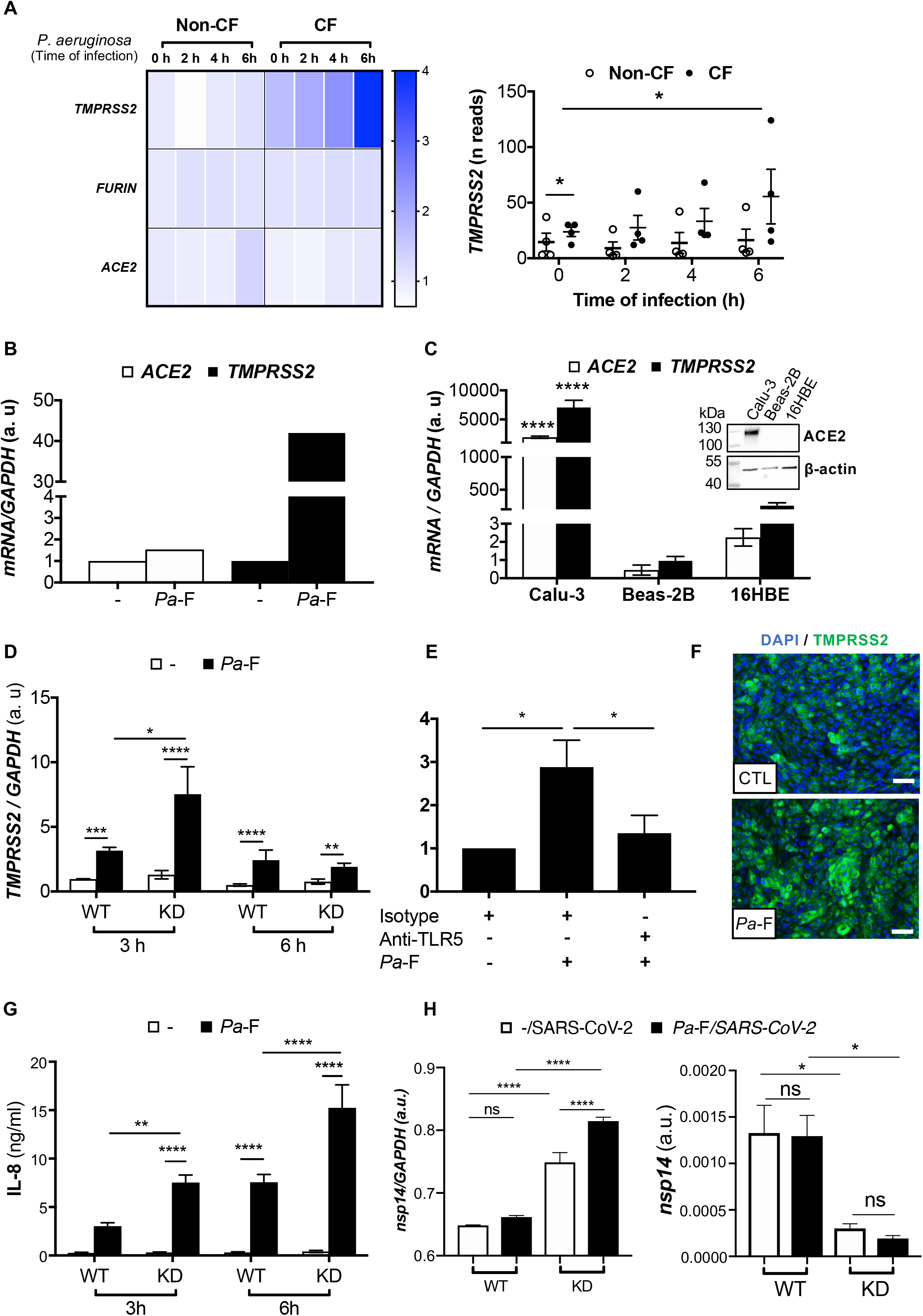
(**A**) (left) Heatmap of *ACE2*, *TMPRSS2* and *FURIN* expression (fold change) and kinetic of (right) *TMPRSS2* expression (reads) in primary hAEC infected by *P. aeruginosa* (RNAseq data extracted from^12^) (non-CF vs. CF at T0, *padjBH=1.76×10^−3^; 0 h vs 6 h in CF, *padjBH=0.048). (**B**) *ACE2* and *TMPRSS2* mRNA expression analysis in submerged CF primary hAEC (n=1) stimulated with control media (reference group) or *Pa*-F (50 ng/ml) for 6 h. (**C)***ACE2* and *TMPRSS2* mRNA expression in submerged cultures of Calu-3, Beas-2B (reference group) and 16HBE14o-cell lines (n=3, ANOVA Dunnett’s multiple comparison test, ***p<0.001). GAPDH is used as housekeeping gene. Western blot (20 μg) of ACE2 protein expression and beta-actin in submerged cultures of Calu-3, Beas-2B and 16HBE14o-cell lines. (**D**) *TMPRSS2* mRNA expression in Calu-3-*CFTR*-WT (reference group) and *CFTR*-KD grown at the air-liquid interface (ALI) and either unstimulated (-) or stimulated 3 h and 6 h by *P. aeruginosa* flagellin (*Pa*-F, 50 ng/ml) (n=5), ANOVA Bonferroni’s multiple comparison test, *p<0.05, ***p<0.001, ****p<0.001). (**E**) Immunofluorescence analysis of TMPRSS2 protein expression (Nucleus are stained with DAPI) in Calu-3 cells grown at the ALI and stimulated by *Pa*-F for 18 h (Scale bar, 40 μM). (**F**) IL-8 production of Calu-3-CFTR-WT and CFTR-KD grown at the ALI and unstimulated or stimulated 3 h and 6 h by *Pa*-F (50 ng/ml) (n=5, ANOVA Bonferroni’s multiple comparison test, *p<0.05, ***p<0.001, ***p<0.0001). (**G**) *TMPRSS2* mRNA expression in Calu-3-*CFTR*-KD grown at the ALI incubated 1h with isotype or anti-TLR5 antibody (10 μg/ml) and then either unstimulated or stimulated 6 h by *Pa*-F (50 ng/ml) (n=3), ANOVA Dunnet’s multiple comparison test, *p<0.05. (**H**) (left) intracellular *nsp14/GAPDH* and (right) apical (supernatants) *nsp14* mRNA expression in Calu-3-*CFTR*-WT (reference group) and *CFTR*-KD grown at the air-liquid interface (ALI) and either unstimulated (-) or stimulated 16 h by *Pa*-F (50 ng/ml), then infected 24 h by SARS-CoV-2 (16) (strain BetaCoV/France/IDF0571/2020 (accession ID EPI_ISL_411218)) (MOI=1) (n=3, ANOVA Bonferroni’s multiple comparison test, *p<0.05, ****p<0.001).

We thus exposed CF primary hAEC to flagellin, the ligand of Toll-Like Receptor (TLR) 5 and the most important proinflammatory factor of *P. aeruginosa* present in the sputum of CF patients (11). We observed that flagellin significantly increases mRNA levels of *TMPRSS2* without increasing those of *ACE2* (**Fig 1B**).

In order to study the mechanism underlying this increase, we used the Calu-3 cell line, which exhibits high *ACE2* and *TMPRSS2* mRNA levels and detectable ACE2 protein expression compared with Beas-2B and 16HBE airway epithelial cell lines (**Fig. 1C**). This result is consistent with the already observed higher ability of SARS-CoV-2 to replicate in Calu-3 cells in comparison to Beas-2B (6).

Next, as observed in CF primary hAEC, we showed that exposure of Calu-3 cells to *PA*-F significantly increases *TMPRSS2* transcripts, without affecting ACE2 mRNA (**Fig. 1D**) and protein expression (not illustrated). Intriguingly, the increase in *TMPRSS2* expression is more important in Calu-3 cells deficient for *CFTR* (Calu-3-*CFTR*-KD) than in standard Calu-3 cells sufficient for *CFTR* (Calu-3-*CFTR*-WT) (**Fig. 1D**). This increase in *TMPRSS2* expression is TLR5 dependent (**Fig. 1E**) and also detected at the protein level (**Fig. 1F**). As expected, flagellin induces the synthesis of the proinflammatory cytokine IL-8 (**Fig. 1G**) both in *CFTR*-WT and -KD Calu-3 cells. As already described (12), CF-like epithelial cells exhibit a more potent inflammatory response to flagellin.

Finally, after infection with SARS-CoV-2, we observed an increase in the intracellular viral mRNA *nsp14*, which is significantly higher in Calu-3-*CFTR*-KD cells and even more important when cells were pre-stimulated with flagellin (**Fig. 1H, left**). In contrast, extracellular viral mRNA *nsp14* level measured at the apical side as a surrogate of viral production is significantly lower in Calu-3-*CFTR*-KD than in Calu-3-*CFTR*-WT cells (**Fig. 1H, right**). Whereas flagellin pre-stimulation does not affect viral particle release in Calu-3-*CFTR*-WT cells, a lower, yet not statistically significant, level of nsp14 mRNA is measured at the apical side of whereas, Calu-3-*CFTR*-KD cells.

The increase of intracellular viral mRNA found in CF cells exposed to flagellin, indicating a higher level of infection, is likely the result of the increased expression of TMPRSS2. Indeed, TMPRSS2 inhibition by the serine protease inhibitor camostat mesylate is sufficient to prevent infection with SARS-CoV-2 (6). We could hypothesize that the lower level of viral particle at the apical side observed in CF cells may be the result of increased host defense capacities of CF cells against viral infection. Further mechanistic works are necessary to know if this response would be protective or damaging for CF patients. Most of CF patients are colonized by *P. aeruginosa* and, to date, even if the results of the clinical follow-up of CF patients are limited, it seems that these patients are not at higher risk to severe COVID-19 than the general population (3). Several studies have delineated the antiviral capacities of flagellin against other respiratory virus, including influenza A (13), and flagellin was recently suggested to modulate the innate response in order to eliminate SARS-CoV-2 and resolve COVID-19 (14). Whether CF patients are protected of developing severe form of COVID-19 thanks to their *P. aeruginosa* infection will be difficult to prove at the clinical level. However, further works elucidating the mechanisms associated with the specific host response of CF cells to SARS-CoV-2 infection could help to elucidate this question and give future therapeutic insights.

## Acknowledgements

LG received a grant from the Faculté de Médecine Sorbonne Université (AAP COVID19). We acknowledge Pr. Marc Chanson (University of Geneva) for generously providing Calu-3-CFTR-WT and Calu-3-CFTR-KD cells (15). We acknowledge Pr. Dieter Gruenert and Dr. Beate Illek (University of California San Francisco (UCSF) for generously providing 16HBE14o-.

## Author Contributions

LG, VB, MR and JB designed experiments, MR, JB, CC, VB, OT, AP, JM and LG made the experiments. LG wrote the manuscript. MR, JB, VB, MRC and HC revised the manuscript.

## Ethics

This project is approved (Opinion number 20-688) by the Inserm Institutional Review Board (IRB00003888, IORG0003254, FWA00005831).

## Supplementary information

### Materials and Methods

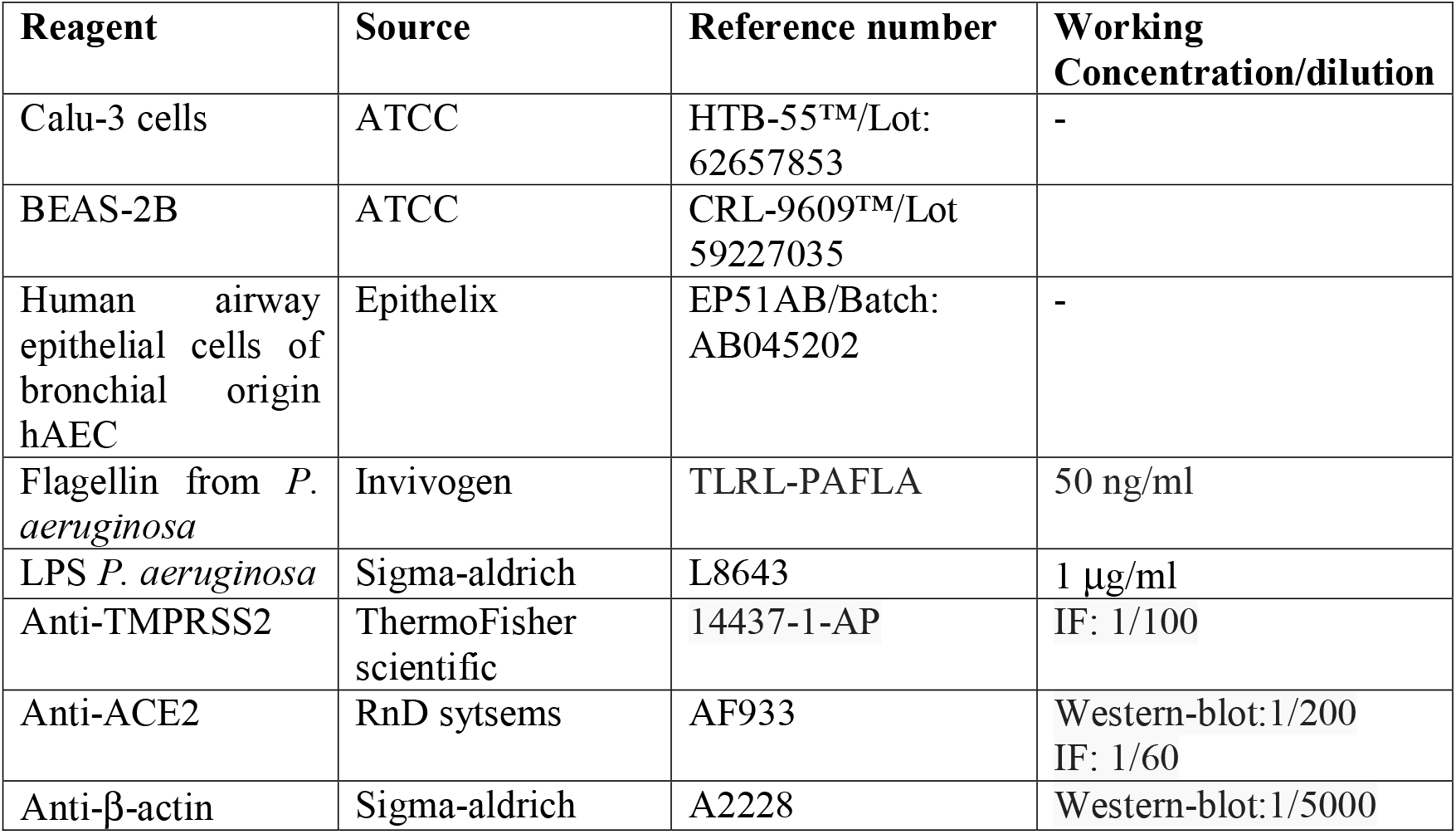
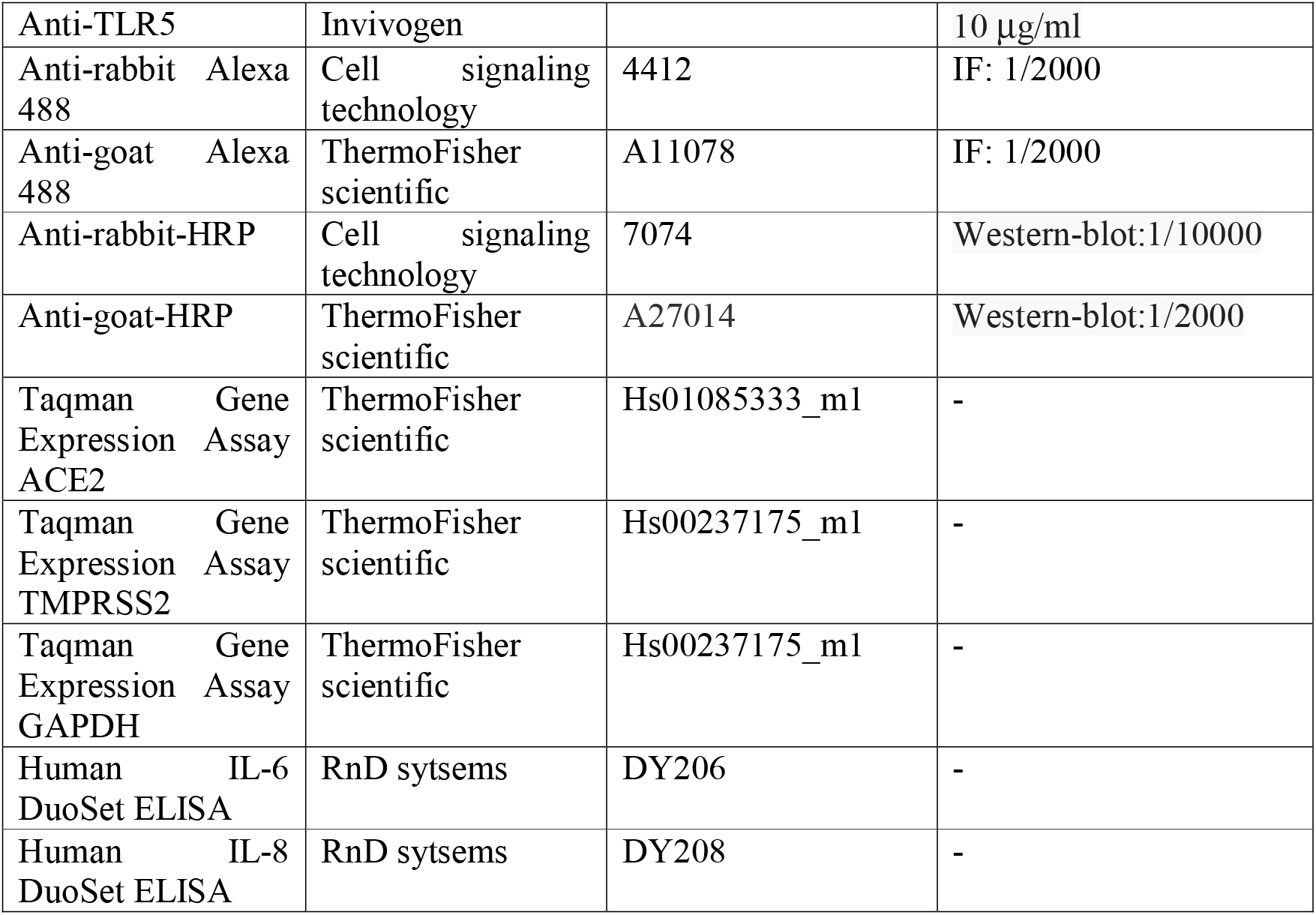

### Cell culture

Calu-3 (ATCC), Calu-3-*CFTR*-WT and Calu-3-*CFTR*-KD (generously given by Pr. M. Chanson, University of Geneva, Switzerland) cells were cultured in MEM-Glutamax (ThermoFisher scientific) supplemented by 10% FCS (Eurobio, Les Ulis, France), 1% non-essential amino acids (ThermoFisher scientific), 10 mM HEPES (ThermoFisher scientific), 1% sodium pyruvate (ThermoFisher scientific) and 1% antibiotics (ThermoFisher scientific) as previously described [16]. Primary CF hAEC (from a male Caucasian donor aged 32 with CF) were cultured as recommended by the manufacturer using hAEC complete culture media (Epithelix, Geneva, Switzerland). Beas-2B cells were cultured in F12 media supplemented by 10% FCS, 10 mM HEPES, and 1% antibiotics. 16HBE14o-were generously given by Dieter Gruenert (originator) and Dr. Beate Illek (provider) from Unversity of California San Francisco (UCSF). They were cultured in MEM-Glutamax supplemented by 10% FCS, and 1% antibiotics as recommended by the provider.

### RT qPCR

RNA was isolated using a NucleoSpin RNA/miRNA (Macherey Nagel, Duren, Germany). RT was performed using a high-capacity cDNA kit (Applied Biosystems, Foster City, CA, USA). Real-time qPCR was performed with an ABI QS3, using Sensifast Probe Lo-Rox Kit (Bio-technofix, Guibeville, France), TaqMan probes and cDNA as a template. For relative quantification, the amount of target genes was normalized to the expression of GAPDH relative to reference group (specified in the figure legends) used as a calibrator and was calculated using the 2^−ΔΔCt^ method.

### Western Blotting

Total proteins were extracted using RIPA buffer (Euromedex, Souffelweyersheim, France). An equal amount of proteins (20 μg) was reduced, size-separated on 12% stain-free precast SDS-polyacrylamide gels (Biorad, Hercules, CA), and transferred to nitrocellulose membranes using iBlot2 apparatus (ThermoFisher scientific). The membranes were blocked in 5% milk in TBS-Tween 0.1% and incubated with specific primary antibodies O/N at 4°C. Bound antibodies were detected using clarity chemiluminescent substrate (Biorad). Images were recorded with a Fujifilm LAS-3000 bioimaging system (Fujifilm, Stamford, CT, USA).

### Immunofluorescence

Cells were plated in 24-well plate on 12 mm diameter #1.5 coverslips (Marienfeld, Lauda-Königshofen, Germany) or on filters at the air-liquid interface and growth as previously described. After treatments, cells were rinsed with PBS and fixed with ice cold PFA 4% 20 min. The cells were permeabilized 10 min with 0.1% Triton X-100 in PBS; and then, washed with PBS and incubated in saturation solution (PBS+BSA 5%) for 1h. Cells were then incubated overnight at 4 °C with primary antibodies in PBS supplemented with 1% BSA. The following day, the cells were washed 3 × 5 min with PBS and incubated for 1h at room temperature with secondary antibodies, followed by DAPI staining. Coverslips were sealed with ProLong diamond mounting media (ThermoFisher scientific). Fluorescence microscopy was achieved using a Olympus fluorescent microscope.

### ELISA

Concentrations of human IL-8 (R&D, Minneapolis, MN, USA) were measured in cell supernatants using ELISA kit according to the manufacturer’s instructions. The 3,3′,5,5′-tetramethylbenzidine substrate was from Cell signaling technology.

### Statistical analysis

Differences among groups were assessed for statistical significance using Prism 7.00 software (GraphPad Software, La Jolla, CA, USA) as indicated in the Figure Legends. Differences with p < 0.05 were considered statistically significant.

## References

1. To T, Viegi G, Cruz A, Taborda-Barata L, Asher M, Behera D, Bennoor K, Boulet LP, Bousquet J, Camargos P, Conceicao C, Gonzalez Diaz S, El-Sony A, Erhola M, Gaga M, Halpin D, Harding L, Maghlakelidze T, Masjedi MR, Mohammad Y, Nunes E, Pigearias B, Sooronbaev T, Stelmach R, Tsiligianni I, Tuyet Lan LT, Valiulis A, Wang C, Williams S, Yorgancioglu A. A Global Respiratory Perspective on the COVID-19 Pandemic: Commentary and Action Proposals. Eur Respir J 2020.

2. Elborn JS. Cystic fibrosis. Lancet 2016; 388: 2519–2531.

3. Cosgriff R, Ahern S, Bell SC, Brownlee K, Burgel PR, Byrnes C, Corvol H, Cheng SY, Elbert A, Faro A, Goss CH, Gulmans V, Marshall BC, McKone E, Middleton PG, Ruseckaite R, Stephenson AL, Carr SB. A multinational report to characterise SARS-CoV-2 infection in people with cystic fibrosis. J Cyst Fibros 2020; 19: 355–358.

4. Chua RL, Lukassen S, Trump S, Hennig BP, Wendisch D, Pott F, Debnath O, Thurmann L, Kurth F, Volker MT, Kazmierski J, Timmermann B, Twardziok S, Schneider S, Machleidt F, Muller-Redetzky H, Maier M, Krannich A, Schmidt S, Balzer F, Liebig J, Loske J, Suttorp N, Eils J, Ishaque N, Liebert UG, von Kalle C, Hocke A, Witzenrath M, Goffinet C, Drosten C, Laudi S, Lehmann I, Conrad C, Sander LE, Eils R. COVID-19 severity correlates with airway epithelium-immune cell interactions identified by single-cell analysis. Nat Biotechnol 2020.

5. Coutard B, Valle C, de Lamballerie X, Canard B, Seidah NG, Decroly E. The spike glycoprotein of the new coronavirus 2019-nCoV contains a furin-like cleavage site absent in CoV of the same clade. Antiviral Res 2020; 176: 104742.

6. Hoffmann M, Kleine-Weber H, Schroeder S, Kruger N, Herrler T, Erichsen S, Schiergens TS, Herrler G, Wu NH, Nitsche A, Muller MA, Drosten C, Pohlmann S. SARS-CoV-2 Cell Entry Depends on ACE2 and TMPRSS2 and Is Blocked by a Clinically Proven Protease Inhibitor. Cell 2020.

7. Walls AC, Park YJ, Tortorici MA, Wall A, McGuire AT, Veesler D. Structure, Function, and Antigenicity of the SARS-CoV-2 Spike Glycoprotein. Cell 2020.

8. Yan R, Zhang Y, Li Y, Xia L, Guo Y, Zhou Q. Structural basis for the recognition of the SARS-CoV-2 by full-length human ACE2. Science 2020.

9. Zhou P, Yang XL, Wang XG, Hu B, Zhang L, Zhang W, Si HR, Zhu Y, Li B, Huang CL, Chen HD, Chen J, Luo Y, Guo H, Jiang RD, Liu MQ, Chen Y, Shen XR, Wang X, Zheng XS, Zhao K, Chen QJ, Deng F, Liu LL, Yan B, Zhan FX, Wang YY, Xiao GF, Shi ZL. A pneumonia outbreak associated with a new coronavirus of probable bat origin. Nature 2020; 579: 270–273.

10. Balloy V, Varet H, Dillies MA, Proux C, Jagla B, Coppee JY, Tabary O, Corvol H, Chignard M, Guillot L. Normal and Cystic Fibrosis Human Bronchial Epithelial Cells Infected with Pseudomonas aeruginosa Exhibit Distinct Gene Activation Patterns. PLoS One 2015; 10: e0140979.

11. Balloy V, Thevenot G, Bienvenu T, Morand P, Corvol H, Clement A, Ramphal R, Hubert D, Chignard M. Flagellin concentrations in expectorations from cystic fibrosis patients. BMC Pulm Med 2014; 14: 100.

12. Blohmke CJ, Victor RE, Hirschfeld AF, Elias IM, Hancock DG, Lane CR, Davidson AG, Wilcox PG, Smith KD, Overhage J, Hancock RE, Turvey SE. Innate immunity mediated by TLR5 as a novel antiinflammatory target for cystic fibrosis lung disease. J Immunol 2008; 180: 7764–7773.

13. Georgel AF, Cayet D, Pizzorno A, Rosa-Calatrava M, Paget C, Sencio V, Dubuisson J, Trottein F, Sirard JC, Carnoy C. Toll-like receptor 5 agonist flagellin reduces influenza A virus replication independently of type I interferon and interleukin 22 and improves antiviral efficacy of oseltamivir. Antiviral Res 2019; 168: 28–35.

14. Golonka RM, Saha P, Yeoh BS, Chattopadhyay S, Gewirtz AT, Joe B, Vijay-Kumar M. Harnessing innate immunity to eliminate SARS-CoV-2 and ameliorate COVID-19 disease. Physiol Genomics 2020; 52: 217–221.

15. Bellec J, Bacchetta M, Losa D, Anegon I, Chanson M, Nguyen TH. CFTR inactivation by lentiviral vector-mediated RNA interference and CRISPR-Cas9 genome editing in human airway epithelial cells. Curr Gene Ther 2015; 15: 447–459.

16. Pizzorno A, Padey B, Julien T, Trouillet-Assant S, Traversier A, Errazuriz-Cerda E, Fouret J, Dubois J, Gaymard A, Lescure FX, Duliere V, Brun P, Constant S, Poissy J, Lina B, Yazdanpanah Y, Terrier O, Rosa-Calatrava M. Characterization and Treatment of SARS-CoV-2 in Nasal and Bronchial Human Airway Epithelia. Cell Rep Med 2020; 1: 100059.

